# Derived ecological niches of indoor microbes are crucial for asthma symptoms in university dormitories

**DOI:** 10.1101/2020.01.05.893529

**Authors:** Xi Fu, Yanling Li, Yi Meng, Qianqian Yuan, Zefei Zhang, Dan Norbäck, Yiqun Deng, Xin Zhang, Yu Sun

**Affiliations:** Guangdong Provincial Key Laboratory of Protein Function and Regulation in Agricultural Organisms, College of Life Sciences, South China Agricultural University, Guangzhou, Guangdong, 510642, PR China; Key Laboratory of Zoonosis of Ministry of Agriculture and Rural Affairs, South China Agricultural University, Guangzhou, Guangdong, 510642, PR China; School of Public Health, Sun Yat-sen University, Guangzhou, PR China; Occupational and Environmental Medicine, Dept. of Medical Science, University Hospital, Uppsala University, 75237 Uppsala, Sweden; Institute of Environmental Science, Shanxi University, Taiyuan, PR China

**Author notes:** These authors contributed equally to this work (Xi Fu, Yanling Li). **Competing interests**: The authors declare that they have no competing interests. **Funding**: We thank for funding support from Guangdong Province (2018KTSCX021), Natural Science Foundation of China (81861138005), Key Research and Development (R&D) Projects of Shanxi Province (201803D31021), Shanxi One Hundred Excellent Experts Project (nr 9) and South China Agricultural University.

## Abstract

Increasing evidences from home environment indicate that microbiome community is associated with asthma. However, indoor microbiome composition can be highly diverse and dynamic, and thus current studies fail to produce consistent association. Chinese university dormitories are special high-density dwellings with a standard built environment and personal characteristics for occupants, which can be used to disentangle the complex interactions between microbes, environmental characteristics and asthma.

Settled air dust and floor dust was collected from 87 dormitory rooms in Shanxi University. Bacterial community was characterized by 16S rRNA amplicon sequencing. Students (n = 357) were surveyed for asthma symptoms.

Asthma symptoms were not associated with the overall bacterial richness, but associated with different phylogenetic classes. Taxa richness and abundance in Clostridia and Bacteroidia were positively associated with asthma (p < 0.05), and these taxa were mainly derived from human gut. Taxa richness (p < 0.1) and abundance (p < 0.05) in Alphaproteobacteria and Actinobacteria were protectively associated with asthma, and these taxa were mainly derived from outdoor environment. Building age, floor and curtain cleaning frequency shaped the overall bacterial community of air dust (p < 0.05). Frequent curtain cleaning increased the relative abundance of 10 protective genera (p < 0.05), and old buildings had mix effects to protective genera (p < 0.05).

Our data shows that taxa from different phylogenetic classes and ecological niches have different health effects, indicating the importance of incorporating evolutionary and ecological concepts in revealing general patterns in the microbiome asthma association analysis.

## Introduction

With the progress of urbanization and industrialization in many developed and developing countries, the prevalence of metabolic and immune diseases has been remarkably increasing in the past few decades, including obesity, diabetes, inflammatory bowel disease, allergies and asthma [1, 2]. Recent culture-independent microbiome studies suggest that the increasing prevalence could be due to loss of microbial diversity in human body [3]. Specifically, the development of asthma is suggested to be associated with reduction and variation of microbiome composition in the outdoor and indoor environment, as well as in the human respiratory and gastrointestinal tract [4–6]. However, it is not easy to identify a consistent pattern between microbial exposure and asthma; many studies produced different or even conflicting results. For example, several studies show that high microbial diversity exposure at home is protective against asthma, especially in the farm environment [7, 8]. But there are several studies indicating that higher microbial diversity does not relate to asthma [9, 10], or even be a risk factor to impair respiratory health [11, 12]. Besides the conflicting results in microbial diversity, the studies surveying the association between relative or absolute abundance of microbial taxa and asthma also produce heterogeneous results. For example, one child cohort study in northeastern United States find only one protective taxa for asthma, whereas another child cohort study in several cities of United States find 373 bacterial taxa are associated with childhood asthma [10, 11].

The first challenge of microbiome analysis is to deal with the complex multidimentional dataset, which contains abundant variations for thousands of taxa. In addition, indoor microbiome composition is affected by many indoor and outdoor environment characteristics, including latitude, climate and humidity, soil and plant, traffic pollution, building design and ventilation, number of occupants and pets, indoor surface material and plumbing systems, settled dust resuspension and so on [13–15]. Microbial community can vary across geographic distance even between rooms in the same building [16–19], and the variation can be temporally changed, as indicated in longitudinal studies in home, hospital and kindergarten rooms [20–22]. The scale of variation could be astonishing even among the same sample type and ecological niche. For example, individual human gut and hand microbiome can be 80%-90% different from one another [23, 24]. Similarly, people inhabit in different locations or different rooms in the same building or different time can be exposed to distinct microbial compositions [18]. Thus, to conduct a clear health association study, either the number of microbiome sampling is large enough to account for the variations, or applying a solid study design to reduce the microbiome variation. Another problem for asthma association analysis is that many personal characteristics are associated with asthma, such as parental asthma, tobacco smoking, obesity, respiratory infection, pollen allergy, age, gender, education level and lifestyle [25], and it is necessary to adjust these variables in the statistical model. However, studies adjusted different set of confounding factors can introduce biases, leading to incomparable conclusions. Overadjustment biases and unnecessary adjustment have also been reported in epidemiologic studies [26]. Overall, the microbiome asthma association study is still in the early exploratory stage, and thus it is optimal to design study with minimal confounding effects to reduce analysis variation.

Chinese university dormitory is a special high-density dwelling. Each room is shared by ~ 4-8 occupants and connected by a long corridor. Rooms within a building share same or similar outdoor characteristics, such as greenness and soil type and traffic pollution, as well as similar indoor characteristics, such as room size and design, window orientation, ventilation system, indoor decoration and furniture. No kitchen is in the dormitory building, and thus no particulate matters emitted from cooking activity. Thus, many factors leading to drastic microbiome variation in home environment are standardized in the dormitory environment. In addition, many confounding variables associated with asthma are also standardized. For example, dormitory occupants have similar age, education level and life style, including eating in the same student canteen, similar sleeping and wake up time and mandatory physical training classes in the university. The participants of this dormitory study are mainly from northern china, which are shown to have low genetic diversity from a previous large-scale genomic survey of China [27].

Few studies have been conducted in Chinese dormitories, and these studies focused on health effect of environmental characteristics, including visible dampness and mold, room ventilation and crowdedness [28, 29]. In this study, we conducted the first microbiome asthma association study in Chinese university dormitories. The analysis aims to disentangle the association between microbial diversity/abundance, environmental characteristics and asthma. Previous hypothesis and observation were also tested in the dormitory dataset.

## Materials and Methods

### Study population

A total of 87 rooms in 10 dormitory buildings were randomly selected in Shanxi University, Taiyuan, China. Photos for building, corridor and dormitory rooms were taken (Figure S1). Settled airborne dust and floor dust were collected for each room in November and December, 2013. Environmental characteristics were recorded. Self-reported questionnaires on asthma related symptoms and personal information were sent to students living in these rooms, and 357 participants responded (response rate 97.3%). The study was approved by University Ethical Committee, and all participants gave their informed consent.

### Assessment of health and environmental data

Questions about current asthma was obtained from the European Community Respiratory Health Study (ECRHS). The asthma related questions included wheeze, breathlessness during wheeze, feeling of chest tightness, shortness of breath during rest, shortness of breath during exercise, woken by attack of shortness of breath [30]. The prevalence of ever had asthma, attack of asthma, and current asthma medication use was very low and not included in the study. Recall time for all these questions were 12 months. The number of asthma symptoms for each participant was counted and categorized as 0, 1, 2, >=3. Questions about current smoking and parental asthma/allergy were also included.

### Dust sampling, DNA extraction and sequencing

Detailed information for dust sampling, DNA extraction and sequencing protocol can refer to a previous publication of microbiome analysis in the same dormitory rooms [31]. Settled airborne and floor dust were collected by petri-dish and vacuum cleaner from 87 randomly selected rooms. A few samples failed to amplify enough DNA for sequencing, and thus in total 83 floor dust and 86 air dust samples were sequenced by Illumina MiSeq platform with MiSeq Reagent Kit v3.

DNA extraction and multiplexed high-throughput sequencing were conducted by Personalbio (www.personalbio.cn). Sequencing data was deposited in Qiita with study ID 12841 (https://qiita.ucsd.edu/study/description/12841).

### Bioinformatics and association analysis

Microbiome data were mainly analyzed by Quantitative Insights Into Microbial Ecology (QIIME, v1.8.0) [32] and R. All samples were rarefied to even depth of 11,000 reads. Beta diversity analysis were conducted by weighted UniFrac distance metrics [33], and visualized by non-metric multidimensional scaling (NMDS) analysis [34].

Bivariate analysis between environmental characteristics and diversity was performed using Kruskal-Wallis test. Environmental characteristics with p values lower than 0.1 were further input in the multivariate analysis by forward stepwise approach in IBM SPSS Statistics (v21.0). Permutational bivariate and multivariate analyses of variance (Adonis) [35] between environmental characteristics and microbiome data were conducted by the vegan package in R.

Associations between bacterial richness/abundance and asthma symptom were examined by hierarchical ordinal regression in STATA (v15.0), adjusted for variables including gender, smoking, parental asthma. Dormitory building was second hierarchy. For each sample, the overall richness of bacteria and richness within the dominant phyla and classes were estimated as number of observed taxonomic units (OTUs) within each phylogenetic lineage. Associations between bacterial abundance and asthma score were analyzed at the phylum, class and genus level. Parallel lines test was conducted to examine the model fitness. If p < 0.01 in the parallel lines test, we further conducted nominal regression. Microbes with significant associations in nominal regression model were kept as associated microbes. The associations between environmental characteristics and protective and risk microbes were examined by linear regression in STATA (v15.0).

## Results

In total, 357 participants were surveyed with health data from 87 randomly selected dormitory rooms. All rooms had natural ventilation by windows and doors. The size of the dormitory rooms ranged from 12 to 23 m^2^ with 4 to 8 occupants in each room, and the average living space was 3.1 m^2^ per person (interquartile range IQR, 2.3-3.5 m^2^). 69.3% of the participants were girls. The mean age of the participants was 21 years (IQR 20-23 years). The prevalence of smoking, parental asthma and pollen allergy was 4.8%, 1.4% and 5.6%, respectively. Attack of shortness of breath during rest and exercise were common (10.4% and 31.7%; Table 1). The prevalence of feeling of chest tight and waken up by attack of shortness of breath were 6.4% and 2.8%. Only 1.1% of participants reported breathlessness during wheeze (Table 1).

**Table 1.**
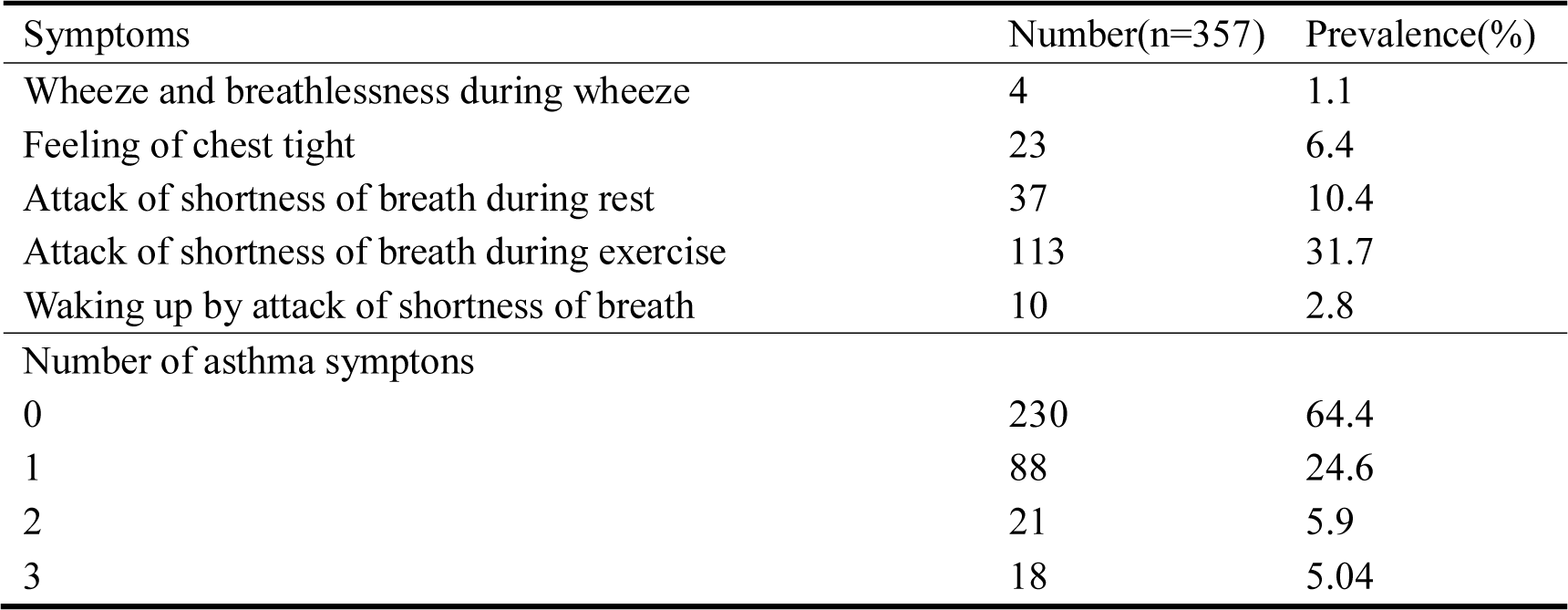
Number and prevalence of asthma symptoms among university students (N = 357) in selected dormitories.

**Table 2.**
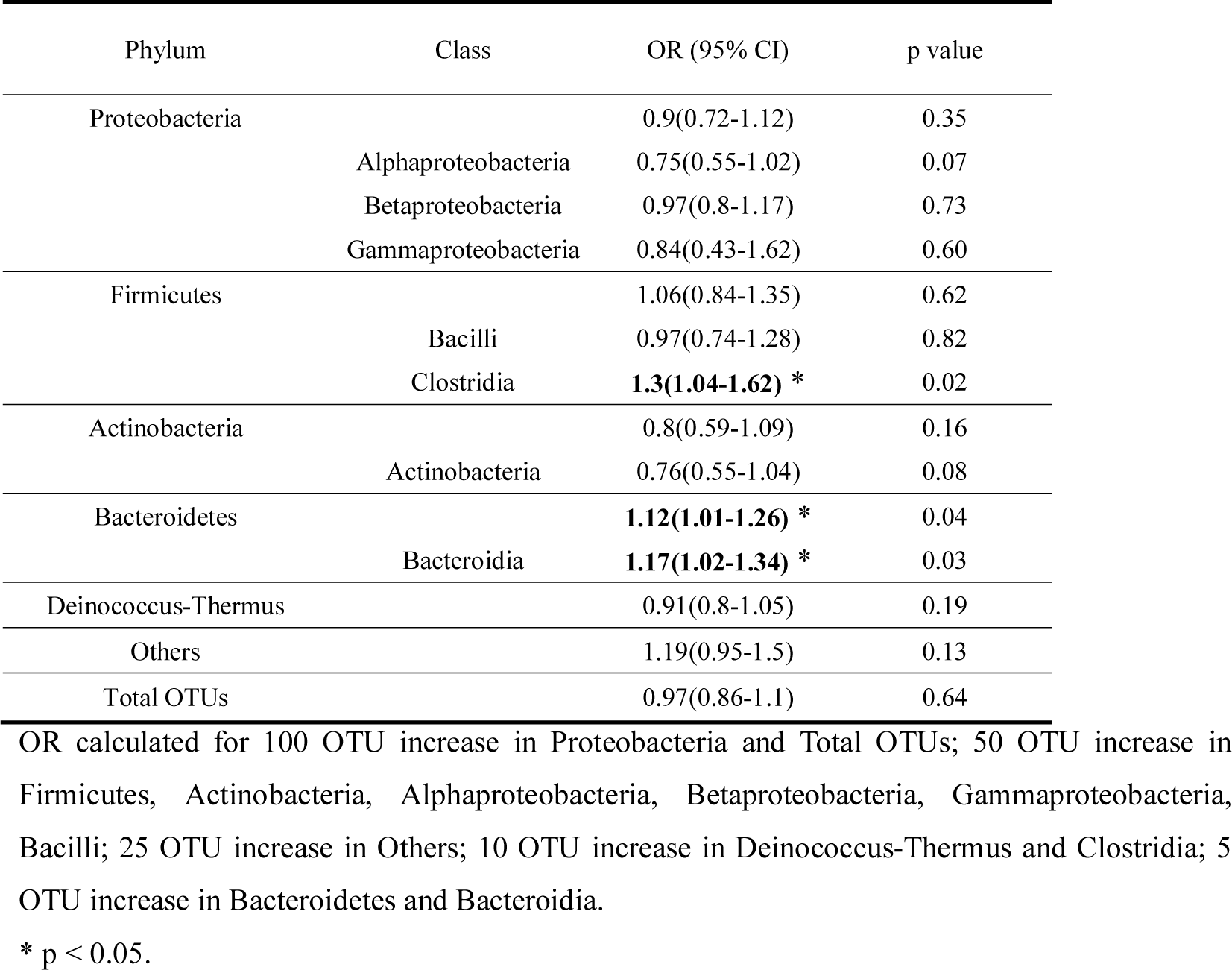
Association between bacterial richness (observed number of OTUs) in air dust and asthma symptoms (N = 357) in dormitories. Odds Ratio (OR) and 95% confident interval (CI) were calculated by 2-level hierarchic ordinal regression models adjusted for gender, smoking and parental asthma. Besides the five major phyla, other phyla had relatively small number of observed OTUs and thus merged as “Others”.

### Association between microbial diversity/abundance and asthma symtoms

Detailed microbiome data analysis in these dormitories has been reported in a previous publication [31]. Overall, the microbial composition of airborne and floor dust were drastically different, as shown in the NMDS (Figure 1) and Adonis analysis (R^2^ = 0.65, p < 0.001). Except two floor samples in building No. 9 that were more similar to the composition of airborne dust, the air and floor dust samples clustered separately along NMDS1. Thus, we conducted association analysis for air and floor dust separately.

**Figure 1.**
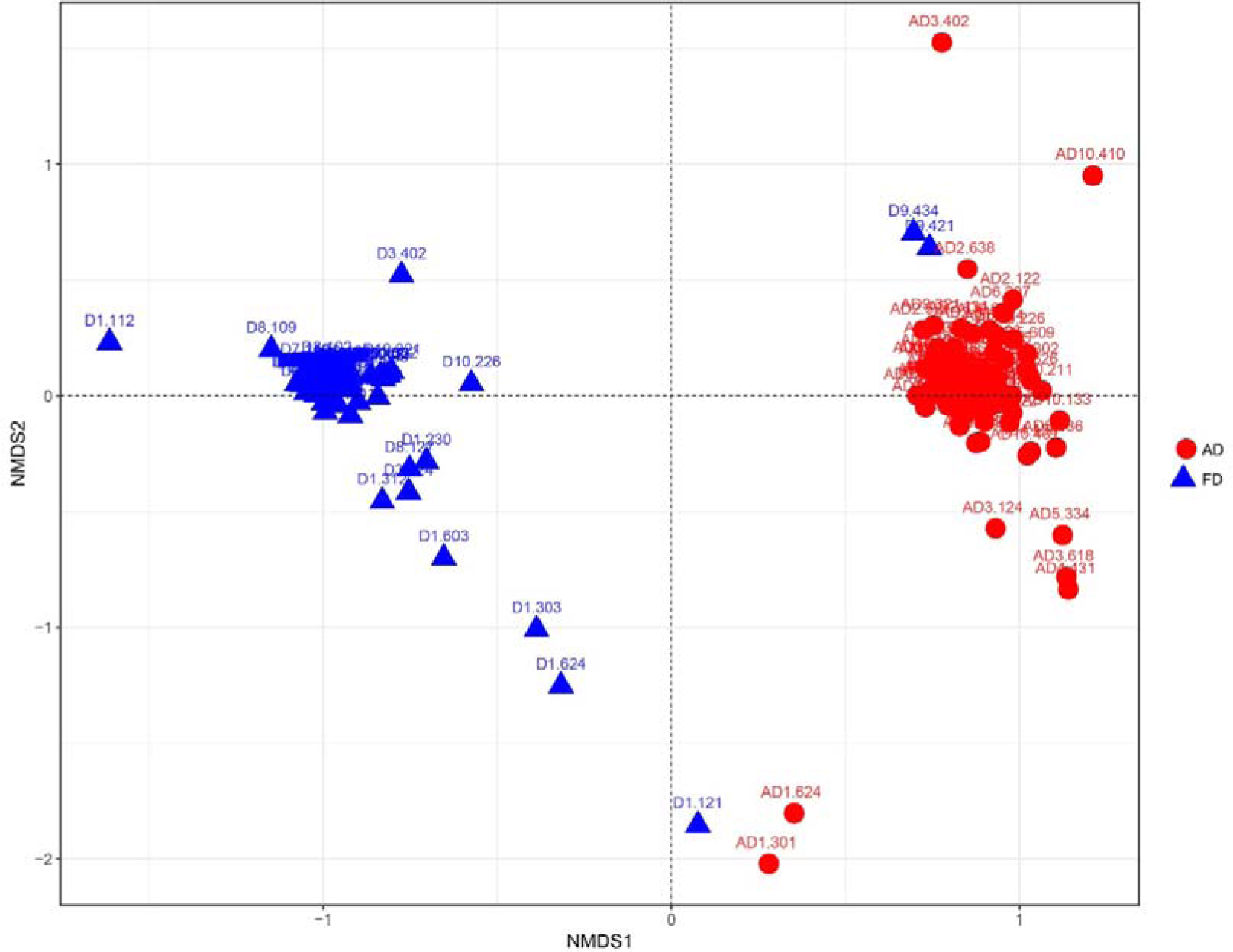
Bacterial community variation between airborne (AD) and floor dust (FD). Non-metric multidimensional scaling (NMDS) analysis of microbial composition based on weighted UniFrac distance was performed. Building and room numbers are labeled for dust samples.

The overall bacterial richness was not associated with asthma symptoms for both floor and air dust (p > 0.2). In air dust, richness in Alphaproteobacteria and Actinobacteria were marginally negatively associated with asthma (p = 0.07 and 0.08), and richness in Clostridia and Bacteroidia were positively associated with asthma (p = 0.02 and 0.03; Table 1). In floor dust, richness in Proteobacteria had a slight trend toward negatively associated with asthma (p = 0.14), and richness in Clostridia had a trend toward positively associated with asthma (p = 0.12; Table S1).

We further calculated the association between relative abundance of each bacterial phylum, class and genus and asthma symptoms (Table S2 and S3). In air dust, we found 5 genera of Alphaproteobacteria and 8 genera of Actinobacteria were protectively associated with asthma (p < 0.05; Table 3), consistent with the richness analysis. Similarly, 4 genera in Clostridia were mainly positively associated with asthma, and *Eubacterium* and *Peptococcus* were also detected in floor dust. Two genera in Bacteroidia were positively associated with asthma. Other classes showed mixed associations with asthma. For example, in the class of Betaproteobacteria, two genera were positively associated and two general were negatively associated with asthma. In floor dust, six genera in the class of Clostridia were positive association with asthma (Table 4), also consistent with the richness analysis.

**Table 3.**
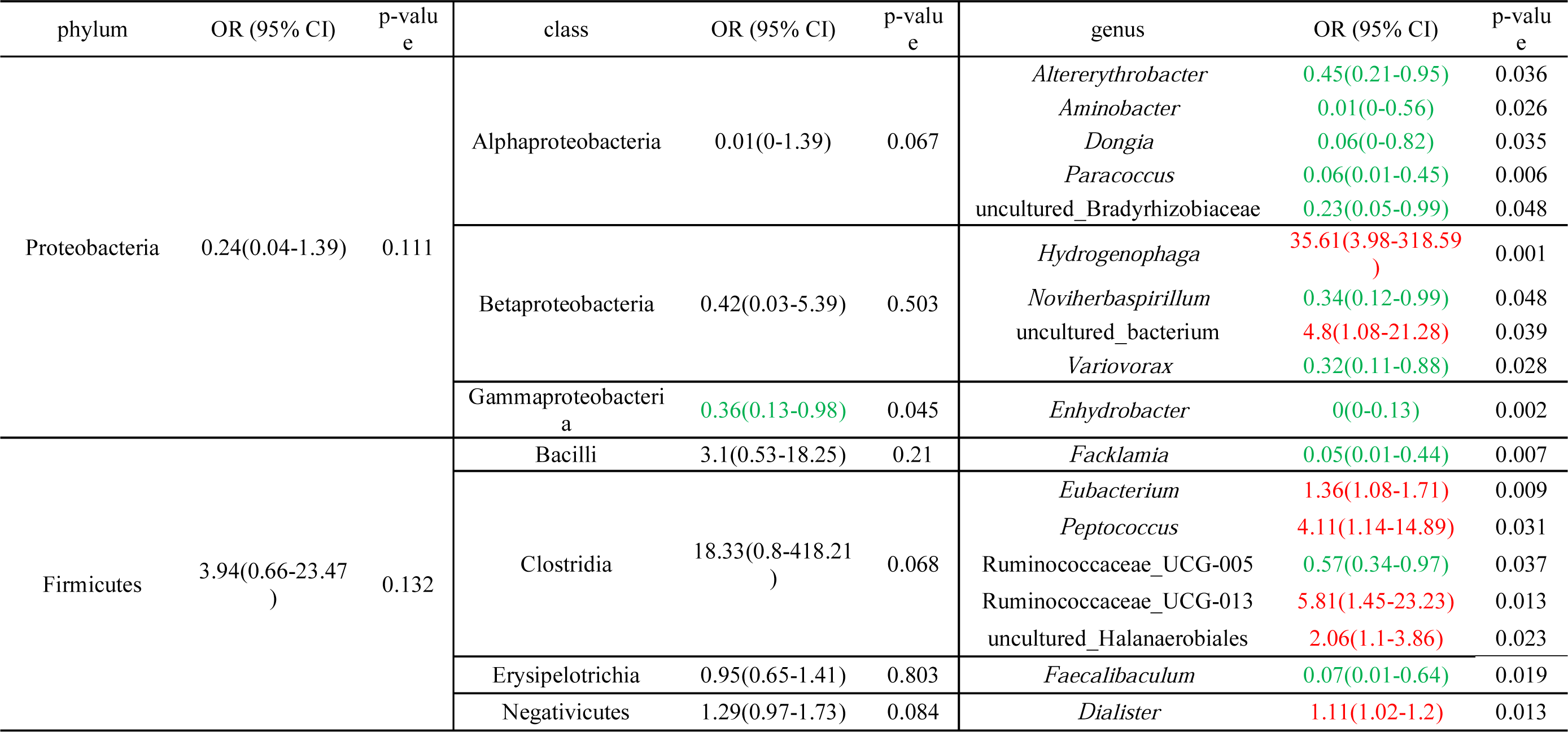
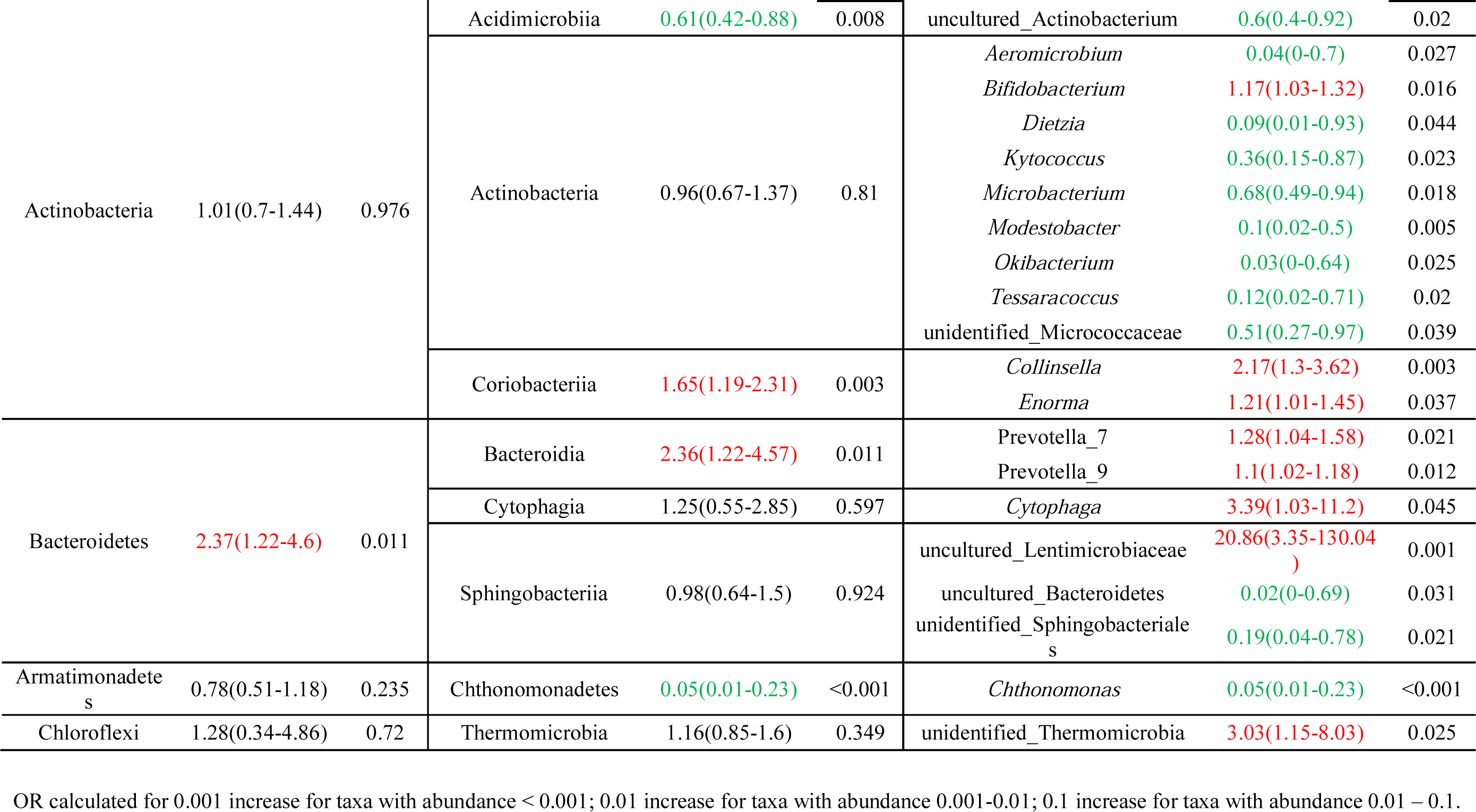
Association between bacterial abundance in air dust and asthma symptoms among student (N = 357) in dormitories. Odds Ratio (OR) and 95% confident interval (CI) were calculated by 2-level hierarchic ordinal regression models adjusted for gender, smoking and parental asthma. Significant positive association is coded with red color and significant negative association is coded with green color.

**Table 4.**
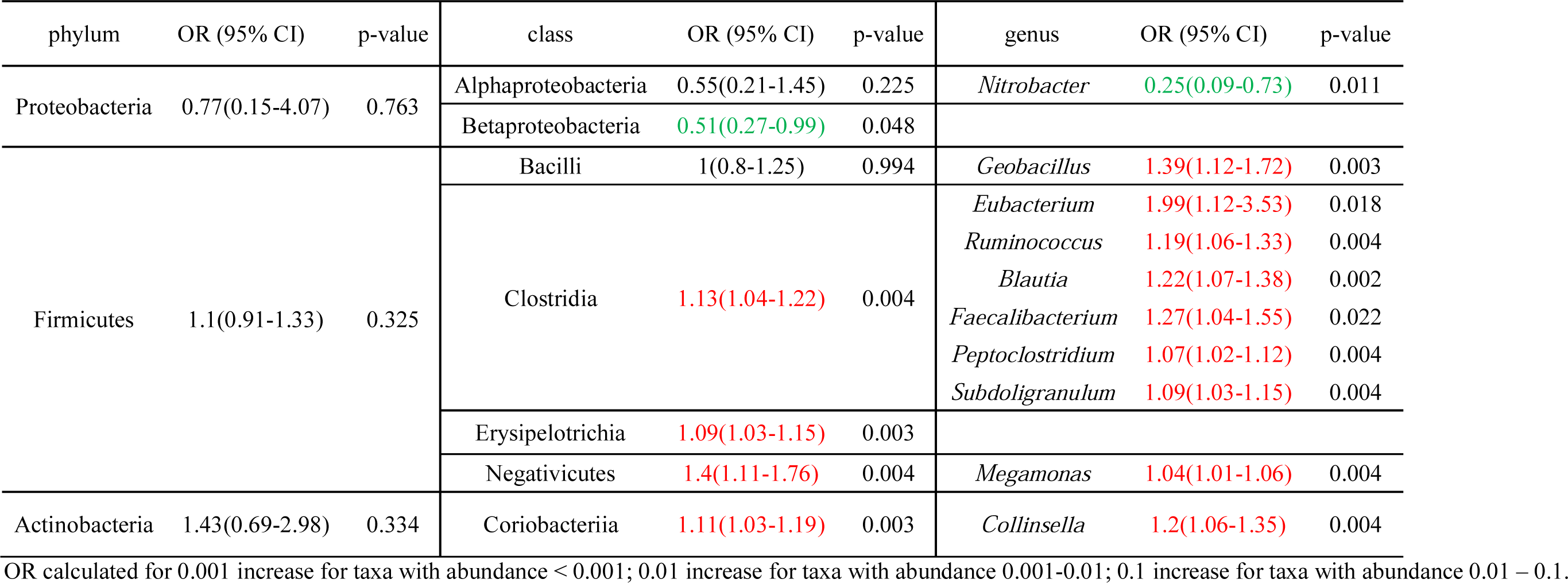
Association between bacterial abundance in floor dust and asthma symptoms among student (N = 357) in dormitories. Odds Ratio (OR) and 95% confident interval (CI) were calculated by 2-level hierarchic logistic regression models adjusted for gender, smoking and parental asthma. Significant positive association is coded with red color and significant negative association is coded with green color (* p < 0.05, ** p < 0.01).

An interesting phenomenon is that taxa from different ecological niches were differently associated with asthma. Alphaproteobacteria and Actinobacteria were mainly derived from outdoor environment, and were presented in low abundance in human derived niches (0.001% and 0.92% in gut, data from Human Microbiome Project [36, 37]), and the protective genera were not detected in human gut. Clostridia and Bacteroidia were dominant classes in human gut. Bacteroidia accounted for 74.0% and Clostridia accounted for 19.4% of human gut microbiota [36, 37]. The risk genera in the indoor environment, including *Prevotella* (3.7%), *Eubacterium* (4.8%), *Ruminococcus* (1.5%), *Blautia* (0.92%), *Faecalibacterium* (3.9%), *Subdoligranulum* (4.3%), *Collinsella* (0.2%) and *Bifidobacterium* (0.63%), were present in high abundance in human gut (abundance values in parenthesis).

A recent indoor microbiome study reported that the sum relative abundance of intercorrelated genera showed much stronger association to asthma than any single genus [38]. Thus, the protective effect can be more accurately assessed by microbial clusters. We conducted factor analysis for the protective and risk microbes in air dust and found a similar pattern (Table S4 and S5). The protective microbes were mainly clustered into 2 factors (Table S4). The first factor included 5 genera of Actinobacteria and 4 genera of Proteobacteria, and the second factor included 3 genera of Actinobacteria and 1 genera of Alphaproteobacteria. Both factors were strongly associated with asthma protection (p = 0.005 and 0.001).

### Association between environmental factors and microbial diversity/abundance

We first conducted the association analysis between environmental characteristics and the overall microbial composition and diversity. Three environmental characteristics were removed in the analysis as they were positively correlated with other characteristics (Table S6). Visible dampness and mold were detected in few rooms, and were also excluded. The bivariate analysis were shown in Table S7 and S8. Selected environmental characteristics (p < 0.1) were further calculated in the multivariate model (Table 5). Building age, curtain cleaning frequency and floor cleaning frequency were associated with bacteria composition in air dust. Building age, sex of occupants and wall surface type were associated with bacterial composition variation in floor dust (p < 0.05). These characteristics can explain 2.7% −4.5% of total variation. East/west window orientation increased bacterial richness in air dust, while plants in room increased bacterial richness in floor dust (p < 0.05).

**Tables 5.** Multivariate analysis between environmental characteristics and bacterial richness (α-diversity) and community variation (β-diversity). The α-diversity was calculated by forward stepwise linear multiple regression, and β-diversity was calculated by forward stepwise Adonis multivariate analysis on 10,000 permutations.

| Air dust |  |  | Floor dust |  |
| --- | --- | --- | --- | --- |
| | observed OTU<br>(α-diversity) | community variation<br>(β-diversity) | observed OTU<br>(α-diversity) | community variation<br>(β-diversity) |
|  | beta (95% CI) | R2 | beta (95% CI) | R2 |
| **Environmental characteristics** |  |  |  |  |
| Building age a |  | 0.035 ** |  | 0.035 * |
| Sex of occupants b |  |  |  | 0.044 ** |
| Floor level c |  |  |  |  |
| Floor surface type d |  |  |  |  |
| Floor cleaning method e |  |  |  |  |
| Floor cleaning frequency f |  | 0.027 * |  |  |
| Wall surface type g |  |  |  | 0.045 ** |
| Window orientation h | 152.18 (38.63, 265.72) ** |  |  |  |
| Curtain cleaning frequency i |  | 0.037 * |  |  |
| Plant in room j |  |  | 40.50 (3.47, 77.53) * |  |

\*\*\* p < 0.001, \*\* p < 0.01, \* p < 0.05
<sup>a</sup> Building age: new (0), old (1); <sup>b</sup> Sex of occupants: male (0), female (1); <sup>c</sup> Floor level: ground (1), top (2), middle (3); <sup>d</sup> Floor surface type: concrete (1), ceramic (2), stone (3), wood (4); <sup>e</sup> Floor cleaning method: dry (1), wet (2); <sup>f</sup> Floor cleaning frequency: < 3 times/week (1), > 3 times/week (2); <sup>g</sup> Wall surface type: paint (1), wall paper (2); <sup>h</sup> Window orientation: south/north (0), east/west (1); <sup>i</sup> Curtain cleaning frequency: 2 times/year (0), 4 times/years (1); <sup>j</sup> Plant in room: no (0), yes (1).

We further conducted association analysis between environmental characteristics and the protective and risk microbes for asthma (Table S9 and S10), and significant associations for air dust samples were summarized in Table 6. Old buildings had higher abundance of several protective bacteria, including *Caulobacter* and 2 unidentified Bacteroidetes, and lower abundance of several protective bacteria, *Microbacterium*, *Tessaracoccus* and an uncultured Bradyrhizobiaceae. Frequent curtain cleaning increased the abundance of 10 protective genera. Other environmental characteristics only affected one asthma-associated microbes.

**Table 6.** Association between environmental characteristics and risk/protective genera in air dust. Beta coefficient was calculated by linear regression models and only significant beta are shown (p < 0.05). Genera identified to be protective in this study are coded with green color, and genera identified to be risk are coded with green color.

|  |  | Building age | Sex of occupants | Floor surface type | Floor cleaning method | Floor cleaning frequency | Wall surface type | Window orientation | Curtain cleaning frequency |
| --- | --- | --- | --- | --- | --- | --- | --- | --- | --- |
| Proteobacteria | *Caulobacter* | 0.13 |  |  |  |  |  |  |  |
|  | Noviherbaspirillum |  |  |  |  | 0.11 |  | -0.64 |  |
|  | Methyloversatilis |  |  |  |  |  |  |  | 0.99 |
|  | uncultured_Bradyrhizobiaceae | -0.11 |  |  |  |  |  |  | 0.22 |
|  | Variovorax |  |  |  |  |  | -0.11 |  | 0.5 |
| Firmicutes | Facklamia |  |  |  | 0.03 |  |  |  |  |
|  | Eisenbergiella |  |  |  |  |  |  |  | 0.2 |
|  | Peptococcus | -0.1 |  |  |  |  |  |  |  |
| Actinobacteria | Microbacterium | -0.43 |  |  |  |  |  |  | 1.63 |
|  | Unidentified_Actinobacteria |  |  |  |  |  |  |  | 0.61 |
|  | Kytococcus |  |  |  |  |  |  |  | 1.04 |
|  | Tessaracoccus | -0.08 |  |  |  |  |  |  |  |
|  | Okibacterium |  |  |  |  |  |  |  | 0.09 |
|  | Dietzia |  |  |  |  |  |  |  | 0.33 |
|  | Unidentified_Acidimicrobiia |  |  |  |  |  |  |  | 1.06 |
| Bacteroidetes | Prevotella_7 |  | -0.5 |  |  |  |  |  |  |
|  | Cytophaga |  |  | -0.03 |  |  |  |  |  |
|  | uncultured_Bacteroidetes | 0.03 |  |  |  |  |  |  |  |
|  | Unidentified_Sphingobacteriia | 0.08 |  |  |  |  |  |  |  |
| Parcubacteria | Candidatus_Sonnebornia_yantaiensis |  |  | 0.07 |  |  |  |  |  |

OR calculated for 0.001 increase for taxa with abundance < 0.001; 0.01 increase for taxa with abundance 0.001-0.01; 0.1 increase for taxa with abundance 0.01 – 0.1.

## Discussion

Here we presented the first microbiome asthma association study in Chinese university dormitories. Similar pattern were observed between the two sampling strategies at higher taxonomic level. For example, bacterial richness and abundance in Clostridia were positively associated with asthma, and Alphaproteobacteria were mainly protectively associated with asthma. But the results were more heterogeneous at lower taxonomic level. We identified 23 protective and 15 risk genera in air dust and 1 protective and 9 risk genera in floor dust, but only 3 risk genera were identified in both dataset. The difference may be partly due to the significant composition variation between air and floor dust, which has been thoroughly analyzed and discussed in a previous paper [31]. One of the major compositional differences was that floor dust had high proportion of *Pseudomonas* (75.1%). The dominant taxon can swamp low frequency organisms in the sequencing and rarefaction analysis, leading to biased microbial richness estimation [39]. Similarly, we found significant lower number of OTUs [31] and asthma-associated taxa in floor dust compared with air dust in dormitories. Thus, we argued that the microbes identified from air dust is more comprehensively represent the associated microbes in this indoor environment, and in the following discussion, we mainly compared the result from air dust with other studies.

In the past few years, the “hygiene hypothesis” and the later expanded version “diversity hypothesis” have drawn a lot of attention and discussion [40]. The hypotheses suggest that the increasing prevalence of metabolic and immune diseases including asthma is related to loss of biodiversity, especially microbial diversity, in human gut and outdoor/indoor environment [1]. The mechanism is suggested to relate to inadequate stimulation and maturation of immunoregulatory circuits in low microbial exposure environment. The theory has gained supports from human gut [1], but heterogeneous results were produced for indoor microbiome studies. A cross-sectional study reported that children lived in farm environment were exposed to greater variety of environmental microorganisms, providing protective effects for asthma and atopy [7], and a few subsequent cohort studies also supported the claim [38, 41]. However, there were several studies failed to identify the pattern [9-11, 42]. Indeed, the number of studies does not support the “diversity hypothesis” is higher than studies support it in the indoor microbiome area. In this study, we found asthma was not associated with the overall bacterial richness, but was associated with specific classes, such as Clostridia, Bacteroidia, Alphaproteobacteria and Actinobacteria. The results need to be further verified in other studies. If the conclusion holds, then the “diversity hypothesis” can be modified as that high exposure in specific classes or lineages, rather than the whole microorganisms, is associated with respiratory health in the indoor environment.

In this study, we found Alphaproteobacteria and Actinobacteria were mainly protective for asthma. Alphaproteobacteria is an extraordinarily diverse and ancient group of bacteria, and its members are widely distributed in various ecological niches, such as soil, water, air and even low-nutrient environments as deep oceanic sediments and glacial ice [43, 44]. Two birth cohort studies showed that Alphaproteobacteria was present in higher abundance in farm homes compared to urban homes, providing protective effect for asthma development [9, 38]. Similarly, Actinobacteria is also a diverse class of bacteria that is widely distributed in terrestrial and aquatic ecosystems [45, 46]. The protective association between Actinobacteria and asthma has been reported in one indoor study [38], and also in a sputum study of nonasthmatic subjects compared with asthmatic subjects [47].

Taxa diversity and abundance in Clostridia were risk factors for asthma in dormitories. Previous studies showed conflicting results between Clostridia and asthma. Studies in gut microbiome reported that colonization of *Clostridium difficile* in youth children was associated with eczema, wheeze, asthma and atopy [48, 49]. Our unpublished study in middle school classrooms of Johor Bahru, Malaysia also reported *Robinsoniella* from Clostridia is a risk factor for asthma. But one case-control study for adults suggested that *Clostridium* cluster XI was protectively associated with asthma prevalence [50]. In addition, *Clostridium butyricum* was suggested to be used as a potential therapeutic microbe for asthma treatment [51]. The discrepancy suggests that Clostridia may have complex effects for asthma. It is possible that the taxa from this class mainly pose risk effects, but exposure with certain genera may be involved in immune training and thus provide protective effects.

An interesting result from this dormitory study is that the protective and risk taxa were mainly derived from different ecological niches. Protective taxa in Alphaproteobacteria and Actinobacteria were mainly from outdoor environment, whereas the risk taxa in Clostridia and Bacteroidia are mainly derived from human gut. Other risk genera, such as *Bifidobacteria* from Actinobacteria and *Collinsella* from Coriobacteriia, also followed the rule and were highly abundant in human gut. Human sheds and releases large amount of microbes to the indoor environment every day. Previous studies reported that students in university classroom emitted ~ 36 million bacterial and 7.3 million fungal genomes per person per hour [52], and the microbiome in a new house can be rapidly changed by new occupants in a few days [21]. A review article summarized that human-derived microbes accounted for 4-40% of the total bacterial load in various indoor environments [53]. Thus, human-derived microbes account for large proportion of indoor microbiome and can further influence the human respiratory health, as shown in this study. The conceptual advances can promote practical advances in the indoor environment. Practical solutions, such as increasing air and microbial exchange between indoor and outdoor environment and frequently cleaning or use less textile fabrics, can promote the respiratory health for indoor occupants.

A strength of this study is that we choose Chinese university dormitory rooms, a standard built environment with similar outdoor and indoor characteristics and relatively low microbiome variation, to disentangle the association between microbial exposure and asthma. The personal characteristics of dormitory occupants were also standardized with similar age, education level and life style. As many potential confounding factors were controlled, consistent results between microbial diversity and abundance analyses were produced, and the protective and risk taxa showed clear phylogenetic pattern with distinct derived ecological niches. A limitation of this study is that due to the small amount of dust collected by petri-dish collector, we cannot conducted the third-generation sequencing with higher taxonomical resolution. Also, the cross-sectional study design limits the possibility to draw conclusions on causal effects.

In conclusion, we present here the first microbiome asthma association study in university dormitories, and identified a list of protective and risk microbes derived from different ecological niches. The study demonstrate the importance of incorporating ecological and evolutionary concepts in disentangle the microbes associated with respiratory health. This study can promote current understanding of microbial exposure and asthma and future practice to maintain healthy indoor microbiome.

## Acknowledgement

We thank Personalbio (www.personalbio.cn) for assistant in sequencing and bioinformatics analysis.

---

**Figure S1.**
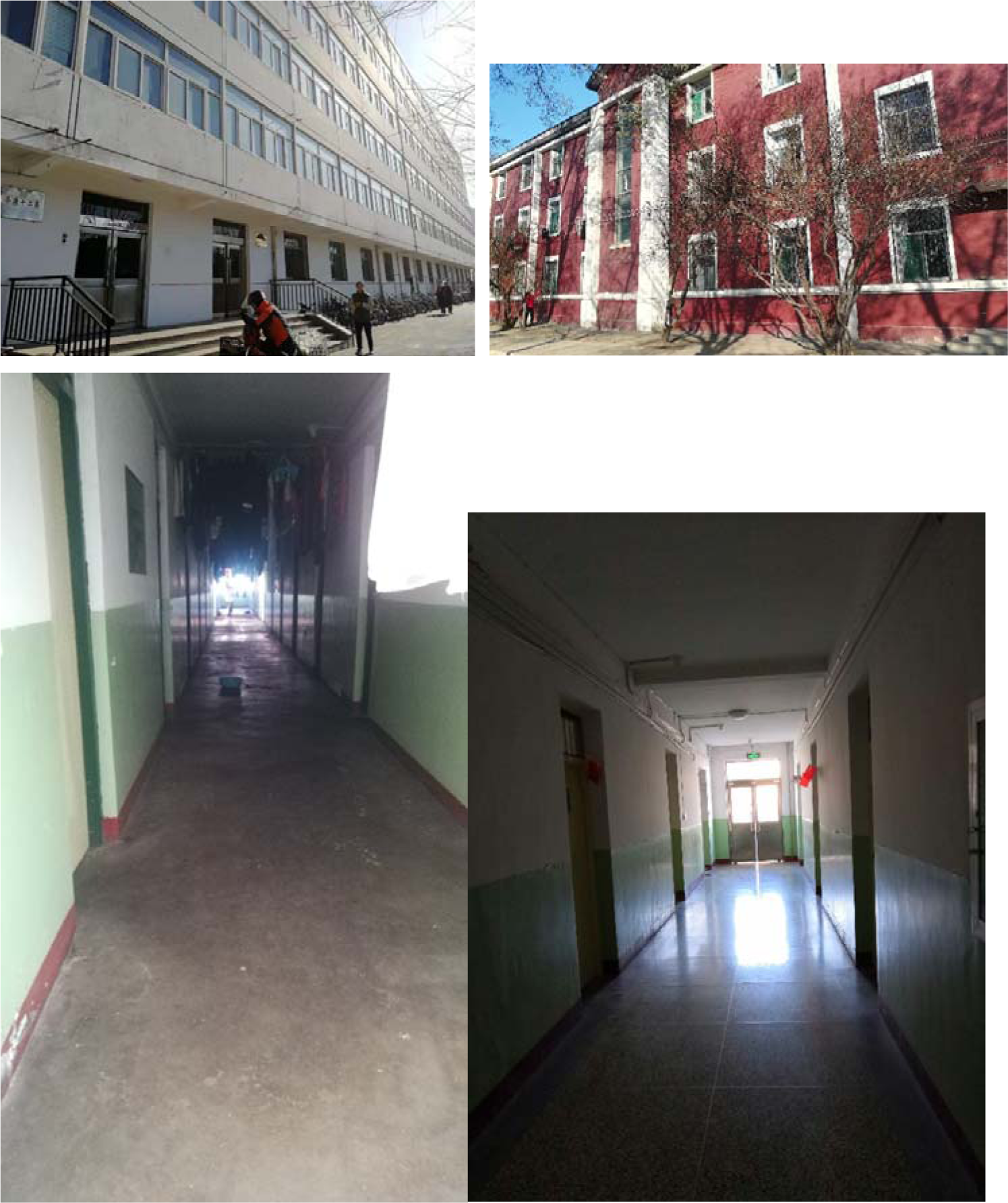
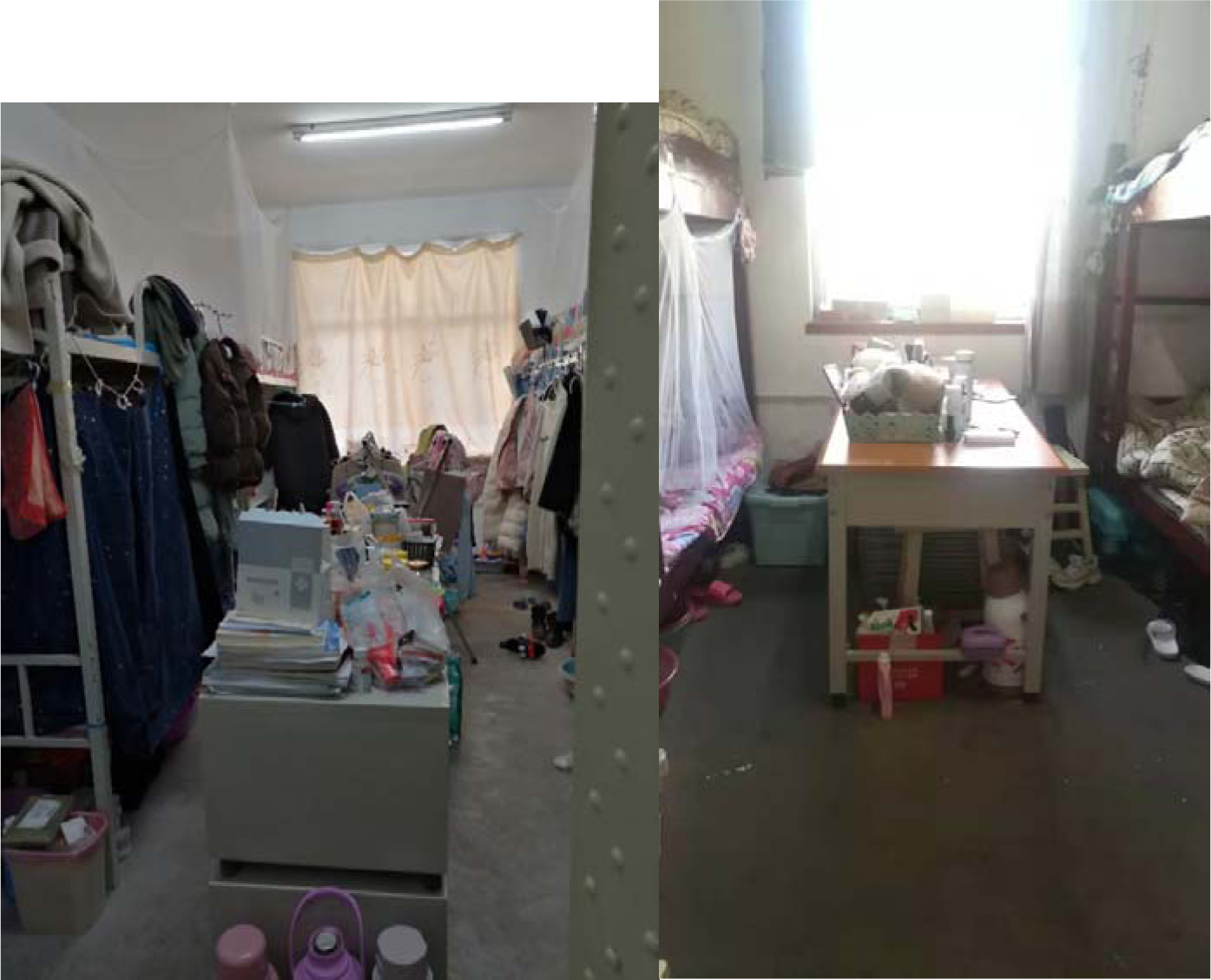
Photos for dormitory building, corridor and rooms.

